# Modeling Functional Genetic Alteration in Cancer Reveals New Candidate Driver Genes

**DOI:** 10.1101/242354

**Authors:** Nadav Brandes, Nathan Linial, Michal Linial

**Affiliations:** School of Computer Science and Engineering, The Hebrew University of Jerusalem, Israel; Department of Biological Chemistry, The Alexander Silberman Institute of Life Sciences, The Hebrew University of Jerusalem, Israel

**Keywords:** Cancer genes, Machine learning, TCGA, ExAC, Positive selection, Census, ClinVar

## Abstract

Compiling the catalogue of genes actively involved in tumorigenesis (known as cancer drivers) is an ongoing endeavor, with profound implications to the understanding of tumorigenesis and treatment of the disease. An abundance of computational methods have been developed to screening the genome for candidate driver genes based on genomic data of somatic mutations in tumors. Most methods rely on detecting genes displaying excessive mutation rates compared to some background model. This approach is susceptible to false discoveries, due to its sensitivity to the assumptions of the background model, such as the need to account for hyper-mutated samples, cancer types and genomic loci. We present a fundamentally different approach. Instead of focusing on the number of mutations, we examine their content, and their expected effects on the functions of genes. We use a machine-learning model to predict functional effect scores of somatic mutations. For each gene, we compare the distribution of observed effect scores with the distribution expected at random, and report genes showing significant bias. By applying our framework on the ~20k protein-coding human genes, we detected 593 genes showing significant bias towards harmful mutations in the context of cancer. In contrast, we found only 6 significant genes biased in the opposite direction. The list of 593 genes, constructed without any prior knowledge of their role in cancer, shows an overwhelming overlap with known cancer driver genes, but also highlights many overlooked genes. These overlooked genes are promising candidates for novel cancer drivers. Our model is generic and is not restricted to the context of cancer. Applying the same framework to data of human-population genetic variation reveals the opposite trend. Unlike cancer, which is dominated by a bias towards harmful mutations, long-term evolution in healthy individuals results a bias towards less harmful mutations. The underlying assumptions of our framework are minimal, making it ideal for analyzing genetic data in search of genes subjected to positive or negative selection. It is fully open sourced and available for installation and use. Our framework presents a substantial development towards the application of state-of-the-art machine-learning algorithms in genetic studies.

## 1. Introduction

Cancer is a genetic disease, dominated by somatic genetic mutations altering key cellular processes such as DNA repair and cell cycle, which lead to genomic instability and high mutation rate of cancerous cells^1,2^. Most arising somatic mutations are considered passenger mutations, whereas only a small fraction of them have a direct role in oncogenesis, and are therefore referred to as cancer driver mutations^3-6^. The study of cancer drivers can be conducted in various resolutions: i) single genetic variants (i.e. driver mutations)^7^, ii) clusters of variants in specific regions of genes^8^, iii) whole genes/proteins (i.e. driver genes)^9^, and iv) pathway networks and cellular processes^10-12^. Most experimental and computational efforts have focused on whole genes and proteins involved in induction and progression of tumors.

Most driver genes are protein-coding^13,14^. Therefore, most studies focus on somatic mutations in coding regions^15^. Most somatic mutations in coding regions are non-synonymous (i.e. missense or nonsense), whereas germline mutations that have fixed during human evolution tend to be synonymous^16,17^.

Compiling a comprehensive catalogue of cancer driver genes is of utmost importance for the study and treatment of cancer. This effort led to the Cancer Gene Census project^14,17^, a comprehensive catalogue of the currently known genes causally implicated in cancer. Other catalogues combining experimental validation and literature support for driver candidates include projects like Candidate Cancer Gene Database (CCGD)^18^. In recent years, cancer genomic research has benefited from ever increasing quantities (and quality) of human molecular data. The Cancer Genome Atlas (TCGA) is a flagship project that provides the most comprehensive resource of cancer genetic data, currently covering >10,000 tumor samples in over 30 cancer types^19^. In the presence of such rich data, the expansion of these catalogues in an ongoing process^20,21^.

Numerous computational frameworks have been designed for the purpose of identifying genes suspected as drivers^9,22,23^. Most of these frameworks are based on the premise that driver genes are recurrent across samples, and are therefore recognized by excessive numbers of somatic mutations. In contrast, passenger mutations are expected to appear at random. This computational approach has identified many key drivers, but it neglects the specific properties of somatic mutations, most notably their functional impact. It is to be expected that driver genes will not only show higher number of mutations, but also mutations that are, on average, more harmful to protein function^24^. Furthermore, assessing whether a gene shows an excessive number of mutations must be considered in view of a null background model. Since cancer is characterized by order-of-magnitudes variability in mutation rate among cancer types^22^, samples and genomic loci, the resulted candidate drivers are highly sensitive to modeling choices, leading to inconclusiveness and controversy^9^.

We propose an alternative conceptual approach that takes into account the exact properties of each coding mutation. Instead of counting mutations, we suggest to evaluate their functional impact, seeking genes with significant bias towards damaging mutations. We define genes with excess of damaging mutations to be “alteration promoting”, which are, in the context of cancer genomics, promising driver candidates. In contrast, long-term human evolution should mostly foster “alteration rejecting” genes. A gene will be said to be showing “alteration bias” if it is either alteration promoting or alteration rejecting.

In this study, we introduce a novel statistical framework for identifying genes with alteration bias. Our framework relies on a machine-learning prediction model that assigns effect scores to mutations in coding regions. We formulated a per-gene background model for the distribution of effect scores expected at random. By comparing this null background distribution with the effect scores of the observed mutations, we can detect genes demonstrating significant functional alteration bias, i.e. alteration promoting or alteration rejecting genes. We named our novel framework FABRIC, standing for Functional Alteration Bias Recovery In Coding-regions.

We applied our framework on two datasets. First, we extracted from TCGA’s public dataset a list of almost 3M somatic mutations across the coding regions of many samples and cancer types. Applying our framework on this dataset revealed genes showing alteration bias in the context of cancer. These are strong candidates for driver genes. We also applied the same pipeline on a second dataset of almost 5M germline variants in the human population, extracted from The Exome Aggregation Consortium (ExAC)^25^. We used this dataset in order to validate our framework, and for revealing genes under evolutionary selection in the human population.

## 2. Results

### 2.1 statistical framework for detecting alteration promoting and alteration rejecting genes

We designed a computational framework to systematically scan for genes showing alteration bias in the context of cancer and in general (Fig. 1). We analyze each protein-coding gene independently, extracting all the SNPs within the coding regions of that gene (Fig. 1a). We then use a pre-trained machine-learning model to assign functional effect scores to each SNP (Fig. 1b), which measure the predicted effects of those variants on the protein function. Synonymous mutations are assigned the score of 1 (gene retains full function), nonsense mutations are assigned with 0 score (gene retains no function), and missense mutations are fed through the machine-learning model to obtain a score between 0 to 1 (intuitively, this number can be thought of as the probability of the protein to retain its function given the mutation). Importantly, this machine-learning model was trained in advance on an independent dataset (extracted from ClinVar^26^; see Online Methods), prior to its application in this framework.

**Fig. 1.**
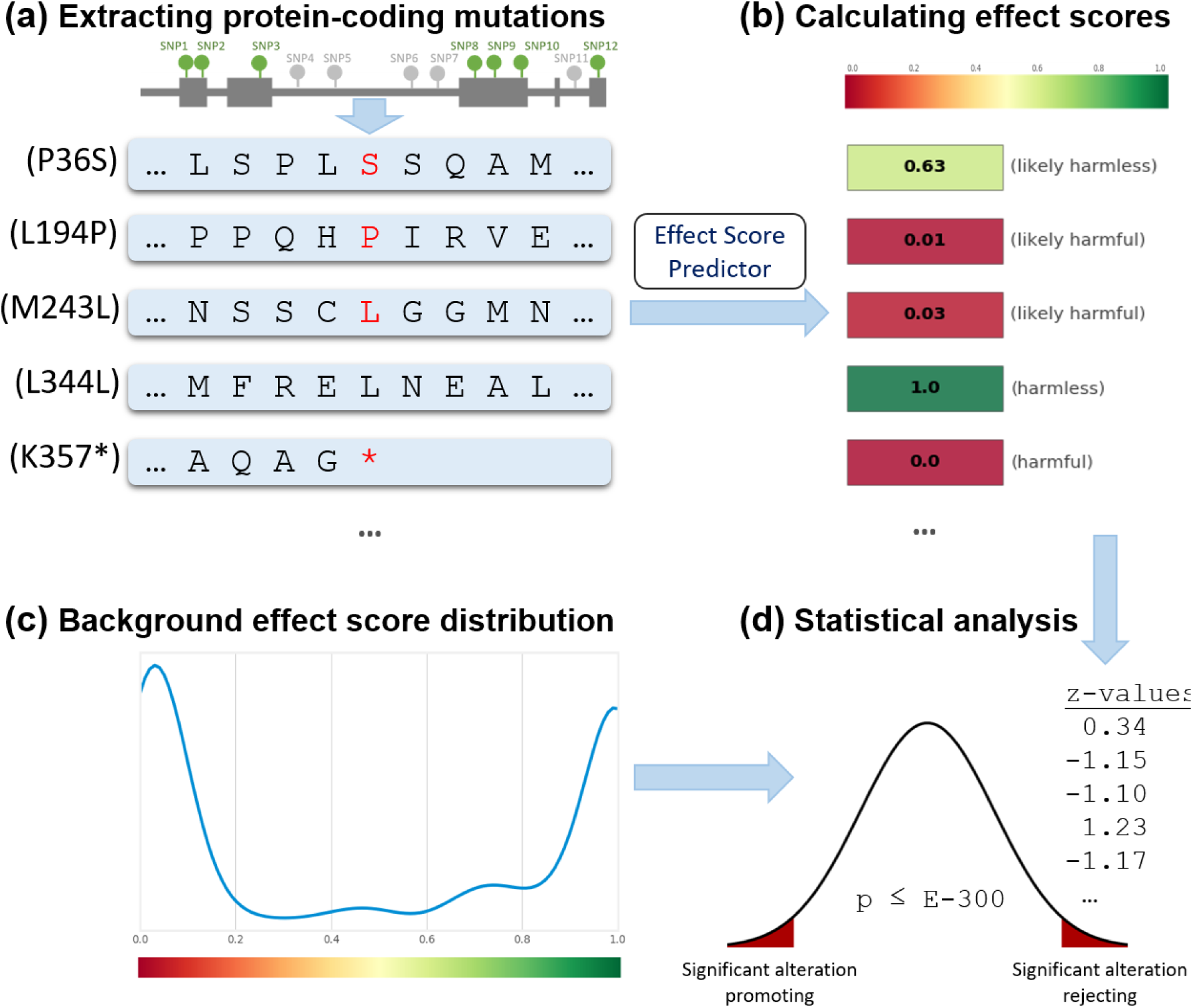
Framework overview. (**a**) All somatic mutations within a particular gene are collected from a variety of different samples and, potentially, different cancer types. SNPs within protein-coding regions are analyzed to study their effects on the protein sequence (synonymous, missense or nonsense), while all other mutations (indels, or mutations outside coding regions) are discarded. (**b**) Using a pre-trained machine-learning model, we assign each mutation a score for its effect on the protein function (lower scores indicate mutations that are more likely harmful). (**c**) In parallel, we consider all possible mutations that could have affected the gene, and assign each of them a similar score, using the same model, in order to construct a precise null background score distribution. The background model is specific for each gene, and also takes into account the frequency of each of the 12 possible nucleotide substitutions in the observed mutations. (**d**) By comparing the observed scores in (b) to their expected distribution calculated in (c), we can calculate z-values for the mutation and a p-value for each gene. Genes with scores that are significantly lower than expected indicate the promotion of harmful alterations, and are suspected as cancer driver genes.

Independently to the calculation of scores for the observed mutations, we construct a background distribution for the expected scores, given that mutations occur at random by a uniform distribution across the gene (Fig. 1c). This background model is precise, and calculated individually for each gene. We then detect significant deviations between the null background distributions to the observed effect scores (Fig. 1d). We calculate z-values to measure the strength of deviation between the observed and expected scores, and, using routine statistical tools (Online Methods), we derive exact p-values. If the average z-value of a gene is significantly negative, it means the observed scores are significantly lower than expected. This indicates they are more harmful to the gene function than expected by the same number of mutations randomly distributed along the gene’s coding sequence. In such case, the gene is deemed to be alteration promoting. Symmetrically, significant positive z-values indicate alteration rejecting genes.

It should be noted that our use of the terms “harmful” and “harmless” in this work refer solely to gene and protein functions at the molecular level, and not to clinical outcomes. If a gene’s function is damaged by some mutation, we will regard this mutation as harmful (at the molecular level), whether or not the function of the gene has clinical significance (at the organism’s level).

The main analysis in this work was conducted on a dataset of somatic mutations in cancer obtained from TCGA^19^, comprised of 3,175,929 somatic mutations across 10,182 samples in 33 cancer types (see Online Methods). From the entire dataset we extracted 2,956,550 SNPs; 2,235,884 of them were in coding regions, and resulted 2,238,945 gene effects. The number of gene effects is slightly higher than the number of coding-region SNPs as, in rare cases, the same mutation may affect overlapping genes.

### 2.2 Precise per-gene background model

Primary concern in the development of this framework was designing a background model that is simple and straightforward, and also strict and robust enough to avoid false discoveries. In particular, a robust statistical background model needs to absorb noise and possible errors of the effect score predictor, which is machine-learning based. The key idea is to use the exact same pre-trained predictor for the extraction of effect scores both in the observed mutations and in the background model. Additionally, it is important to account for the exact number of observed mutations, and their tendency towards specific nucleotide substitutions. Accounting for these factors is critical, as it is well established that mutation rate and nucleotide substitution profiles vary substantially among cancer types, samples and genes^27,28^. We achieve this by considering the effects of all possible mutations within each gene. Notably, our approach ensures that our analysis is insensitive to gene lengths.

For illustration purposes, let us focus on a particular gene: TP53, one of the most studied cancer driver genes^29^. Let us examine the analysis details of this particular gene, as a way to demonstrate the detection of genes showing alteration bias by our framework in general. Of the ~3M somatic mutations extracted from TCGA, 3,167 SNPs were in coding regions of the TP53 gene. These SNPs are categorized as follows: 92 synonymous mutations, 2,563 missense mutations, and 512 nonsense mutations. Synonymous and nonsense mutations are always assigned an effect score of 1 and 0, respectively. The average score of the 2,563 missense mutations in TP53 was 0.02. Altogether, the average score of the 3,167 mutations observed in TP53 was 0.05.

In order to tell whether an average score of 0.05 is significantly low (indicating an alteration promoting gene), it has to be compared against some background model. The designed background model is straightforward (Fig. 2). First, the empirical single-nucleotide substitution frequencies of the observed SNPs are calculated (Fig. 2a). Each of the 4 DNA nucleotides can be substituted to each of the other 3, resulting a matrix with 12 entries summing up to 100%.

**Fig. 2.**
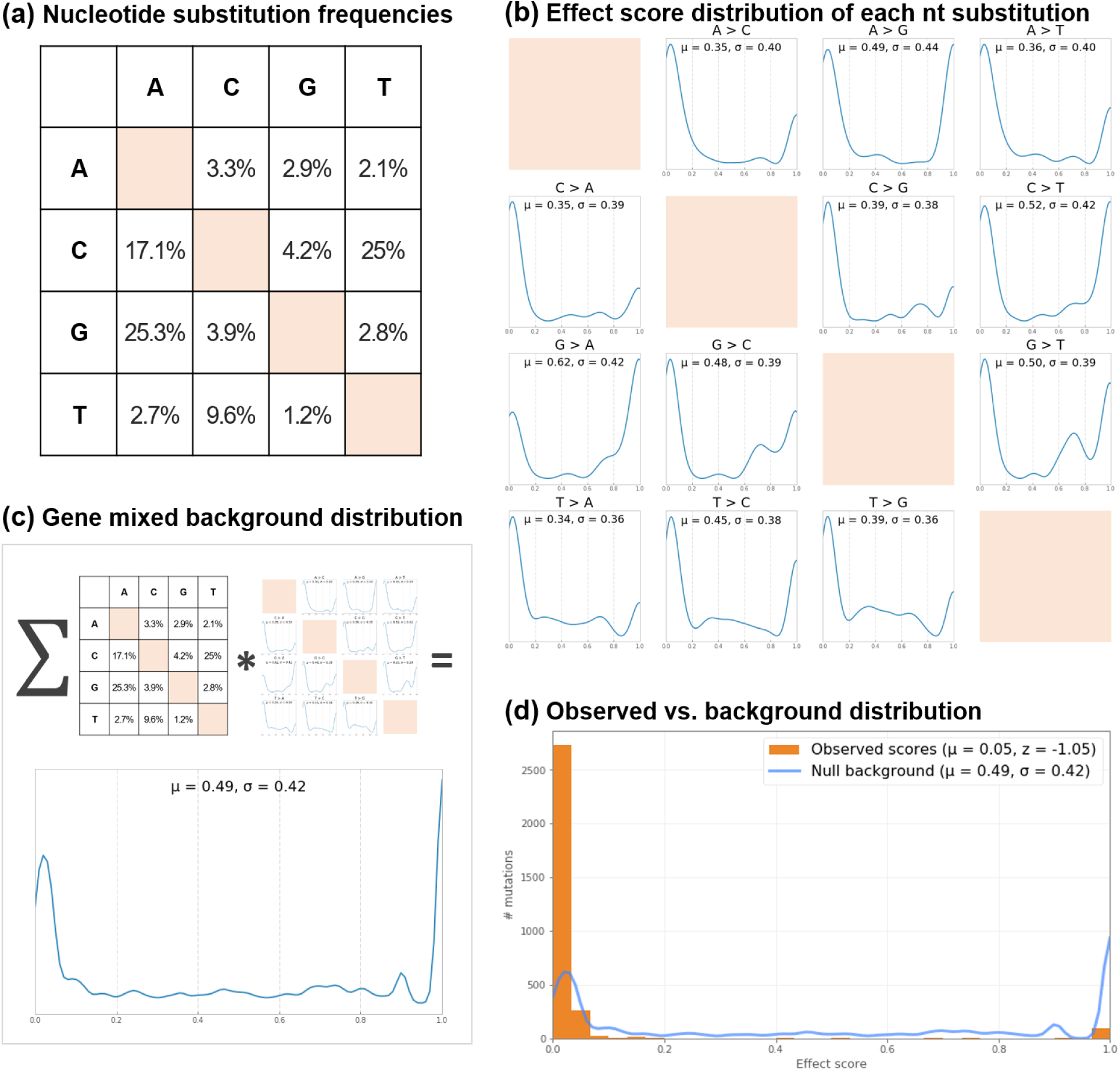
Background model (TP53 as an example). (**a**) Our final dataset contains 3,167 observed SNPs in coding regions of the TP53 gene. A 4x4 matrix of single-nucleotide substitution frequencies was calculated based on these observed mutations. Note that this matrix is very non-symmetric (e.g. among the 3,167 observed mutations in TP53, 25.3% of the substitutions are G>A, but only 2.9% are A>G). (**b**) For each of the 12 possible nucleotide substitutions, an independent background effect score distribution was calculated, by considering all possible substitutions within the coding region of the gene and processing them with our effect score prediction model (the same one used to calculate the scores of the observed mutations). (**c**) By mixing these 12 distributions with the weights of the substitution frequencies, we obtained the gene’s final score distribution, used as the null background model for that specific gene. (**d**) According to the null background distribution, we would expect mutations within the TP53 gene to have a mean score of 0.49. However, the observed mean score of the 3,167 mutations is 0.05, 1.05 standard deviations below the mean (p-value < E-300).

Independently to the calculation of this matrix, for each of the 12 possible substitutions, a background effect score distribution is computed by considering all possible SNPs of that substitution (Fig. 2b). For example, the A-to-G substitution (denoted A>G) has 233 possible distinct occurrences in the sequence of the TP53 gene (because the A nucleotide appears exactly 233 times in the coding region of that gene). Of these, 92 substitutions to G will result a synonymous mutation (assigned with an effect score of 1), 141 will result a missense mutation, and 0 will result a nonsense mutation. The mean score of the 141 missense mutations, calculated by the same predictor, is 0.16. Overall we get that if an A>G substitution occurred in random within the coding region of the TP53 gene, the expected value of the effect score would be 0.49. A similar calculation reveals that the standard deviation of the background distribution of the A>G substitution is 0.44 (the full distribution is plotted in Fig. 2b). An identical calculation is made for all other 11 possible substitutions within the gene.

The final null background distribution, against which the effect scores of the observed mutations is compared, is a simple mixture of these 12 substitution-specific distributions, weighted by their observed frequencies, as calculated in the first step (Fig. 2c; see Online Methods for the mathematical formulation). The resulted mixed score distribution for the TP53 gene has an expected value of 0.49 and standard deviation of 0.42. Intuitively, the background model described the effect score distribution we would expect to see if mutations within the gene arose by a uniform random distribution across its coding sequence, given the observed tendency of mutations towards specific nucleotide substitution frequencies. Finally, the scores of the observed mutations within the gene are compared against its null distribution (Fig. 2d). It appears that a mean score of 0.05 is indeed very low, with overwhelming significance (p-value < E-300), indicating that the TP53 gene is a major alteration promoting gene in cancer, as we would expect from such a dominant oncogene.

### 2.3 A catalogue of alteration promoting genes in cancer

Our analysis, illustrated above for the TP53 gene, was conducted over the entire set of coding human genes, independently for each gene. Of the 17,828 analyzed genes (which contained at least one mutation), the somatic mutations in 593 genes were significantly more harmful than expected at random (FDR q-value < 0.05). The full results of all analyzed genes is available in Supplementary Table S1-TCGA_combined. A short excerpt with the top 20 results is given in Table 1 (sorted by significance according to q-values). It appears that significant alteration promoting genes can dramatically vary in their total number and density of mutations. For example, TP53, the most significant alteration promoting gene in our analysis, has 2.69 SNP mutations per coding-region nucleotide in our dataset, while KMT2D, which is also highly significant (q-value = 5.9E-72), has a 38-fold lower mutation density (0.07 SNP mutations per coding-region nucleotide). Even though TP53 is the most significant alteration promoting gene (with respect to the calculated q-value), the effect score z-values of APC (-1.31) and ARID1A (-1.47) are even lower than that of TP53 (-1.05), indicating a potentially stronger effect size.

In contrast to the abundance of alteration promoting genes, we found only 6 alteration rejecting genes in the context of cancer somatic mutations. Those 6 genes (MUC5B, COL5A1, CDIP1, TSHZ2, TMEM150B and TSHZ1) also show much less impressive statistical significance (lowest FDR q-value is 0.0075).

**Table 1.** Top 20 alteration promoting genes.

| Gene Symbol | Gene Name | Chr | # Observed mutations | Mutation density per nt | Expected mean score | Observed mean score | Score z-value | FDR q-value | Census annotations<sup>a</sup> |
| --- | --- | --- | --- | --- | --- | --- | --- | --- | --- |
| TP53 | tumor protein p53 | 17 | 3,167 | 2.69 | 0.49 | 0.05 | -1.05 | 0 | oncogene, TSG, fusion |
| PIK3CA | phosphatidylinositol-4,5-bisphosphate 3-kinase catalytic subunit alpha | 3 | 1,508 | 0.47 | 0.39 | 0.15 | -0.68 | 4E-240 | oncogene |
| APC | APC, WNT signaling pathway regulator | 5 | 997 | 0.12 | 0.84 | 0.49 | -1.31 | 7.3E-215 | TSG |
| KRAS | KRAS proto-oncogene, GTPase | 12 | 783 | 1.38 | 0.24 | 0.04 | -0.61 | 1.5E-177 | oncogene |
| ARID1A | AT-rich interaction domain 1A | 1 | 640 | 0.09 | 0.82 | 0.45 | -1.47 | 3.6E-169 | TSG, fusion |
| BRAF | B-Raf proto-oncogene, serine/threonine kinase | 7 | 819 | 0.36 | 0.41 | 0.10 | -0.77 | 3.5E-145 | oncogene, fusion |
| PTEN | phosphatase and tensin homolog | 10 | 656 | 0.54 | 0.33 | 0.06 | -0.70 | 9.5E-116 | TSG |
| IDH1 | isocitrate dehydrogenase (NADP(+)) 1, cytosolic | 2 | 536 | 0.43 | 0.33 | 0.05 | -0.68 | 4.8E-94 | oncogene |
| CDKN2A | cyclin dependent kinase inhibitor 2A | 9 | 294 | 0.63 | 0.55 | 0.19 | -0.99 | 9.9E-82 | TSG |
| FBXW7 | F-box and WD repeat domain containing 7 | 4 | 442 | 0.21 | 0.52 | 0.20 | -0.86 | 8.8E-81 | TSG |
| KMT2D | lysine methyltransferase 2D | 12 | 1,191 | 0.07 | 0.88 | 0.72 | -0.65 | 5.9E-72 | oncogene, TSG |
| NF1 | neurofibromin 1 | 17 | 697 | 0.08 | 0.72 | 0.52 | -0.67 | 6.9E-56 | TSG, fusion |
| NRAS | neuroblastoma RAS viral oncogene homolog | 1 | 286 | 0.50 | 0.34 | 0.06 | -0.72 | 3.3E-52 | oncogene |
| RB1 | RB transcriptional corepressor 1 | 13 | 335 | 0.12 | 0.54 | 0.28 | -0.82 | 1.2E-50 | TSG |
| CTNNB1 | catenin beta 1 | 3 | 442 | 0.19 | 0.53 | 0.30 | -0.68 | 6.7E-49 | oncogene, fusion |
| SMAD4 | SMAD family member 4 | 18 | 249 | 0.15 | 0.46 | 0.17 | -0.80 | 3.9E-41 | TSG |
| FAT1 | FAT atypical cadherin 1 | 4 | 1,118 | 0.08 | 0.88 | 0.78 | -0.47 | 7.4E-36 | TSG |
| KMT2C | lysine methyltransferase 2C | 7 | 1,078 | 0.07 | 0.86 | 0.75 | -0.46 | 9.6E-35 | TSG |
| PBRM1 | polybromo 1 | 3 | 346 | 0.07 | 0.68 | 0.47 | -0.74 | 1.6E-34 | TSG |
| EP300 | E1A binding protein p300 | 22 | 496 | 0.07 | 0.75 | 0.56 | -0.61 | 5.6E-33 | TSG, fusion |

a – TSG, Tumor Suppressor Gene

### 2.4 Dissecting the signal for alteration bias

Our analysis gathers information from two distinct signals: i) the type of mutations (synonymous, missense or nonsense) and, ii) the predicted effect scores of missense mutations. As synonymous and nonsense mutations are assigned constant effect scores (1 and 0, respectively), and only the scores of missense mutations vary, these two components are fully orthogonal.

To determine the contribution of each of the two complementary components to our overall analysis, we re-analyzed each gene by these two components separately. Analyzing genes only by the types of mutations, ignoring the varying impacts of missense mutations, was performed by a chi-squared test for the deviation of the observed mutation type frequencies from the frequencies expected by the background model (see Online Methods). Similarly, to analyze genes only by missense effect scores, we simply ignored the other two types of mutations (synonymous and nonsense). In this setting, the background model was adjusted accordingly to disregard synonymous and nonsense mutations as well.

While our main overall analysis found 593 significant alteration promoting genes, the mutation-type and missense analyses found only 387 and 492 significant alteration promoting genes, respectively. The full per-gene results of these analyses are also listed in Supplementary Table S1-TCGA_combined.

As these two complementary analyses are completely orthogonal, no overlap between them should be expected unless for some biological signal. Indeed, we found a significant overlap between them (p-value = 7.96E-30). The number of alteration promoting genes overlapping by the two methods (62 genes) is 2.5 times greater than the number expected at random (24 genes). This non-random signature hints towards the involvement of evolutionary forces in the makeup of cancer somatic mutations which are captured by each of the methods.

### 2.5 Cancer alteration promoting genes strongly overlap with known drivers

The list of 593 significant alteration promoting genes includes strong candidates for cancer driver genes. Indeed, a substantial and significant overlap is found between this list to external lists of cancer driver genes (Fig. 3). We compared our results against three prominent resources of cancer driver genes: Census^13,14^, CCGD^18^ and the MutSig framework^22,30,31^. In CCGD, we considered genes supported by at least 10 published studies. Of the 17,828 genes in our analysis, these three resources identify 526, 366 and 106 genes as cancer drivers, respectively. Our overall analysis (the left bar in each of the six panels, Fig. 3) shows an outstanding overlap with known cancer drivers (enrichment of x6.74, x5.83 and x20.42 with Census, CCGD and MutSig, respectively; p-value ≤ 2.2E-34). This indicates that alteration promoting genes are indeed very promising candidates for cancer drivers.

**Fig. 3.**
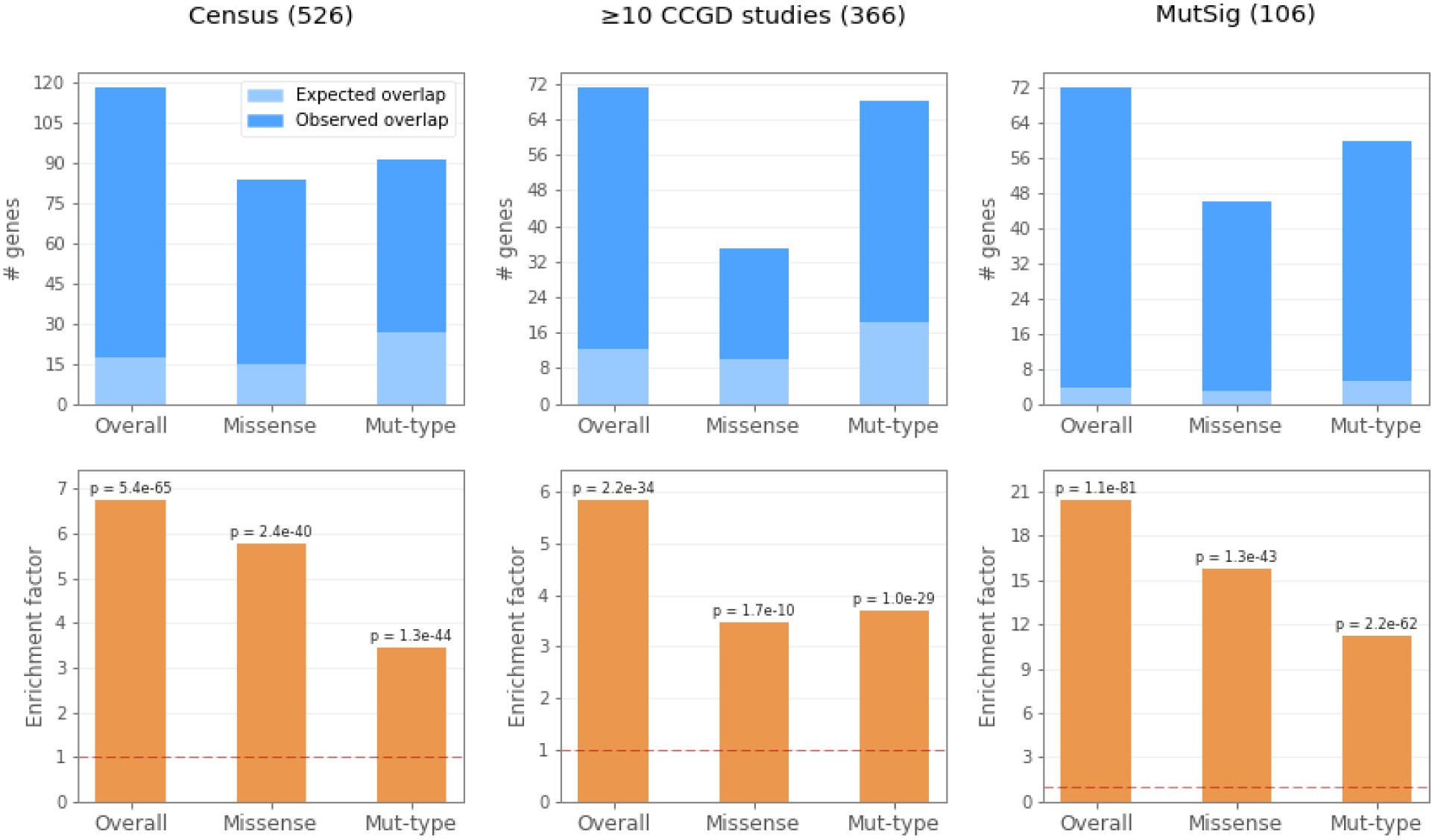
Alteration promoting genes substantially overlap with known cancer drivers. We compared the list of significant alteration promoting genes obtained by our analysis against three external resources for cancer drivers: Census, CCGD and MutSig (see main text). We show three variations of our analysis, each considering different elements of the data: The mutation-type analysis (abbreviated Mut-Type) considers only deviations in the types of mutations (i.e. synonymous, missense or nonsense), treating all missense mutations the same; the missense analysis considers only missense mutations, looking for significant differences between their observed to expected effect scores, while disregarding synonymous and nonsense mutations altogether; the overall analysis combines both signals, by assigning scores to all mutations. The top panel (blue) shows the total number of overlapping genes between each of our three analyses to each of the three compared resources, and the number of genes that would be expected to overlap at random (given hyper-geometric distribution). Note that the total number of genes passing statistical prerequisite conditions for the Mut-Type analysis is only 7,674 of the overall 17,828 (see Online Methods), which is important for the calculation of expected overlaps. The ratio between the observed to the expected number of overlapping genes is defined as the enrichment factor for each pair, and is shown on the lower panel (orange). The baseline of x1 enrichment expected at random is presented as horizontal dashed red lines. Hyper-geometric p-values for the significance levels of the overlaps are also shown.

Fig. 3 also shows the overlaps of known driver genes with each of the two complementary components described earlier: mutation types and missense effect scores. While the mutation-type and missense analyses are capable of recovering much of the signal, the overall integrated analysis shows superior results (see bottom orange panels). A more exhaustive overlapping analysis is available in Supplementary Table S2. To summarize, we find that both the analysis of mutation types and the analysis of missense effect scores are adequate to recovering driver genes, yet even better results are obtained when the two components are combined together, as in our overall analysis. A particularly remarkable enrichment is observed between our framework to the MutSig suite^31^. 72 of the 106 genes reported by MutSig were independently found by our framework (x20.4 enrichment, p-value = 1.15E-81). This strongly supports the validity of both approaches.

### 2.6 Overlooked candidate driver genes

Of the 593 significant alteration promoting genes, we found an outstandingly large subset to overlap with known drivers (Fig. 3), yet many of the reported genes are novel driver candidates without any signature in the commonly used resources. We define a significant alteration promoting gene to be novel if it does not appear in Census and not supported by any study reported in CCGD. 183 of the 593 significant genes meet these criteria, and are listed in Supplementary Table S1-TCGA_combined_overlooked. The most significant overlooked gene is ranked 54th in the overall list of the 593 alteration promoting genes. In other words, the 53 most significant alteration promoting genes we found had already been reported as drivers according to the aforementioned criteria. Of the list of 183 overlooked alteration promoting genes, 26 were defined as very significant (q-value ≤ 0.001), and are listed in Table 2. As we have already presented strong evidence that alteration promoting genes are promising candidates for cancer drivers, we anticipate that among the most significant overlooked ones, many will be established with a direct role in cancer.

**Table 2.** Very significant overlooked driver candidates.

| Rank | Gene Symbol | Gene Name | Chr | # Observed mutations | Mutation density per nt | Score z-value | FDR q-value |
| --- | --- | --- | --- | --- | --- | --- | --- |
| 54 | GRM5 | glutamate metabotropic receptor 5 | 11 | 441 | 0.12 | -0.31 | 2.8E-08 |
| 61 | USP28 | ubiquitin specific peptidase 28 | 11 | 240 | 0.07 | -0.43 | 1.5E-07 |
| 63 | MICU3 | mitochondrial calcium uptake family member 3 | 8 | 88 | 0.06 | -0.64 | 3.9E-07 |
| 69 | ZNF750 | zinc finger protein 750 | 17 | 174 | 0.08 | -0.51 | 2.1E-06 |
| 71 | PGR | progesterone receptor | 11 | 215 | 0.08 | -0.41 | 2.6E-06 |
| 72 | ZNF14 | zinc finger protein 14 | 19 | 160 | 0.08 | -0.47 | 2.9E-06 |
| 82 | HOXA4 | homeobox A4 | 7 | 55 | 0.06 | -0.81 | 8.7E-06 |
| 84 | WTIP | Wilms tumor 1 interacting protein | 19 | 46 | 0.04 | -0.84 | 9.6E-06 |
| 93 | CDKN1A | cyclin dependent kinase inhibitor 1A | 6 | 69 | 0.14 | -0.63 | 4.1E-05 |
| 94 | ELF3 | E74 like ETS transcription factor 3 | 1 | 108 | 0.10 | -0.49 | 5.2E-05 |
| 101 | 40603 | membrane associated ring-CH-type finger 11 | 5 | 121 | 0.10 | -0.47 | 1.1E-04 |
| 103 | PDYN | prodynorphin | 20 | 159 | 0.21 | -0.39 | 1.3E-04 |
| 105 | MYH7 | myosin heavy chain 7 | 14 | 724 | 0.12 | -0.18 | 1.4E-04 |
| 111 | BHLHE22 | basic helix-loop-helix family member e22 | 8 | 58 | 0.05 | -0.70 | 2.4E-04 |
| 112 | PIK3R2 | phosphoinositide-3-kinase regulatory subunit 2 | 19 | 132 | 0.06 | -0.41 | 2.5E-04 |
| 118 | HS3ST4 | heparan sulfate-glucosamine 3-sulfotransferase 4 | 16 | 159 | 0.12 | -0.39 | 2.8E-04 |
| 121 | FOXA2 | forkhead box A2 | 20 | 93 | 0.07 | -0.50 | 2.9E-04 |
| 122 | SPTY2D1 | SPT2 chromatin protein domain containing 1 | 11 | 147 | 0.07 | -0.42 | 3E-04 |
| 126 | HCN4 | hyperpolarization activated cyclic nucleotide gated potassium channel 4 | 15 | 273 | 0.08 | -0.30 | 3.8E-04 |
| 129 | SOX17 | SRY-box 17 | 8 | 97 | 0.08 | -0.50 | 4.1E-04 |
| 130 | MBD1 | methyl-CpG binding domain protein 1 | 18 | 162 | 0.09 | -0.38 | 4.3E-04 |
| 134 | KCNA5 | potassium voltage-gated channel subfamily A member 5 | 12 | 317 | 0.17 | -0.27 | 5.5E-05 |
| 138 | UNCX | UNC homeobox | 7 | 45 | 0.03 | -0.78 | 5.9E-04 |
| 145 | UBXN2B | UBX domain protein 2B | 8 | 62 | 0.06 | -0.56 | 7.5E-04 |
| 149 | CPNE4 | copine 4 | 3 | 209 | 0.13 | -0.31 | 8.4E-04 |
| 154 | ZNF627 | zinc finger protein 627 | 19 | 89 | 0.06 | -0.47 | 9.6E-04 |

### 2.7 Alteration promoting genes across cancer types

Most of our analysis, which has been presented until that point, deals with alteration bias from a pan-cancer perspective. In this pan-cancer analysis, all the somatic mutations extracted from TCGA were combined into a single pool (per gene), disregarding from which samples or cancer types they originated. An important benefit of this setting was the acquiring of the needed statistical power for the analysis, obtained by maximizing the number of samples. However, a notable heterogeneity exists among cancer types in the dominance of driver genes^14,32^. In order to also highlight important differences that might exist in alteration promoting genes among different cancer types, we also conducted the same analysis within each cancer type separately. Those analyses involved the calculation of a specific background model for each combination of gene and cancer type, based on the observed mutations in each combination. This approach overcomes biases caused by inherent differences that exist among cancer types, such as cancer-specific nucleotide substitution frequencies^3,27^.

The summary statistics of our analysis for each analyzed gene in each cancer type is available in Supplementary Table S1. Table 3 lists the 33 analyzed cancer types, ranging from ~40 to ~1,000 samples and ~2,000 to ~900,000 somatic mutations in each. Note the substantial heterogeneity in the rate of mutations per sample among and within cancer types.

**Table 3:** Alteration promoting genes across cancer types.

| TCGA Project | Disease | # Samples | # Mutations | Avg. (std) mutations per sample | # Significant Alteration Promoting Genes | # Significant Diff Genes<sup>a</sup> |
| --- | --- | --- | --- | --- | --- | --- |
| BRCA | Breast Invasive Carcinoma | 986 | 120,988 | 122.7 (347.8) | 12 | 6 |
| LUAD | Lung Adenocarcinoma | 567 | 208,180 | 367.2 (386.3) | 14 | 5 |
| UCEC | Uterine Corpus Endometrial Carcinoma | 530 | 886,377 | 1,672.4 (4,159.7) | 146 | 24 |
| HNSC | Head and Neck Squamous Cell Carcinoma | 508 | 102,309 | 201.4 (288.1) | 15 | 7 |
| LGG | Brain Lower Grade Glioma | 508 | 35,556 | 70.0 (635.7) | 9 | 3 |
| PRAD | Prostate Adenocarcinoma | 495 | 29,286 | 59.2 (408.3) | 3 | 0 |
| LUSC | Lung Squamous Cell Carcinoma | 492 | 181,116 | 368.1 (317.6) | 12 | 2 |
| THCA | Thyroid Carcinoma | 492 | 10,899 | 22.2 (52.3) | 4 | 1 |
| SKCM | Skin Cutaneous Melanoma | 467 | 392,571 | 840.6 (1,423.4) | 16 | 11 |
| STAD | Stomach Adenocarcinoma | 437 | 213,144 | 487.7 (929.3) | 10 | 1 |
| OV | Ovarian Serous Cystadenocarcinoma | 436 | 75,168 | 172.4 (178.7) | 3 | 0 |
| BLCA | Bladder Urothelial Carcinoma | 412 | 134,513 | 326.5 (378.2) | 36 | 15 |
| COAD | Colon Adenocarcinoma | 399 | 264,786 | 663.6 (1,360.4) | 19 | 5 |
| GBM | Glioblastoma Multiforme | 393 | 82,765 | 210.6 (964.8) | 9 | 1 |
| LIHC | Liver Hepatocellular Carcinoma | 364 | 54,238 | 149.0 (161.5) | 5 | 0 |
| KIRC | Kidney Renal Clear Cell Carcinoma | 336 | 26,693 | 79.4 (123.3) | 5 | 3 |
| CESC | Cervical Squamous Cell Carcinoma and Endocervical Adenocarcinoma | 289 | 103,405 | 357.8 (1,157.9) | 12 | 3 |
| KIRP | Kidney Renal Papillary Cell Carcinoma | 281 | 23,765 | 84.6 (40.1) | 1 | 0 |
| SARC | Sarcoma | 237 | 28,159 | 118.8 (281.7) | 2 | 0 |
| ESCA | Esophageal Carcinoma | 184 | 45,313 | 246.3 (317.1) | 4 | 0 |
| PCPG | Pheochromocytoma and Paraganglioma | 179 | 2,411 | 13.5 (7.4) | 2 | 0 |
| PAAD | Pancreatic Adenocarcinoma | 178 | 29,959 | 168.3 (1,534.7) | 5 | 0 |
| TGCT | Testicular Germ Cell Tumors | 144 | 3,198 | 22.2 (12.1) | 2 | 0 |
| LAML | Acute Myeloid Leukemia | 143 | 9,905 | 69.3 (271.7) | 6 | 0 |
| READ | Rectum Adenocarcinoma | 137 | 64,804 | 473.0 (1,783.4) | 10 | 2 |
| THYM | Thymoma | 123 | 4,737 | 38.5 (120.6) | 2 | 1 |
| ACC | Adrenocortical Carcinoma | 92 | 10,747 | 116.8 (316.1) | 0 | 0 |
| MESO | Mesothelioma | 82 | 3,827 | 46.7 (43.8) | 3 | 0 |
| UVM | Uveal Melanoma | 80 | 1,856 | 23.2 (58.6) | 3 | 2 |
| KICH | Kidney Chromophobe | 66 | 2,896 | 43.9 (116.5) | 1 | 0 |
| UCS | Uterine Carcinosarcoma | 57 | 10,449 | 183.3 (681.7) | 6 | 0 |
| CHOL | Cholangiocarcinoma | 51 | 5,503 | 107.9 (220.0) | 2 | 0 |
| DLBC | Lymphoid Neoplasm Diffuse Large B-cell Lymphoma | 37 | 6,406 | 173.1 (106.1) | 1 | 0 |

<sup>a</sup>Diff Genes, Genes with significantly different alteration bias compared to other cancer types

It is important to note that like in every statistical analysis, the potential to find significant results is strongly dependent on the number of available observations, even though the density of somatic mutations in not a factor in our statistical analysis. Some cancer types have a very limited number of available samples in TCGA (Table 3), resulting very few, if any, significant alteration promoting genes. This merely reflects an inevitable lack of statistical power, and it does not mean that those cancer types are not dominated by alteration promoting genes. In total, we found 380 cancer-type specific alteration promoting genes, involving 231 unique genes.

To further highlight cancer-type patterns, we plot the magnitude of alteration bias of selected genes across cancer types (Fig. 4). Fig. 4a shows the 40 most significant alteration promoting genes we found (the first 20 are listed in Table 1). Fig. 4b shows the very significant overlooked candidate driver genes (Table 2). We also looked for genes with significant differences among cancer types (Fig. 4c), aiming to capture specific genes within specific cancer types that show a significant alteration bias compared to the same genes in the other cancer types. Note that a significant difference among cancer types does not necessarily imply a significant bias from the background. For example, TP53, ranked at the top of the list of alteration promoting genes (Fig. 4a), shows only a weak difference in alteration bias across cancer types (ranked 65 of 68 genes; Fig. 4c), confirming its universal role as a driver across cancer types. A full analysis of the differences among cancer types of all analyzed genes is available in Supplementary Table S1-TCGA_diff.

**Fig. 4.**
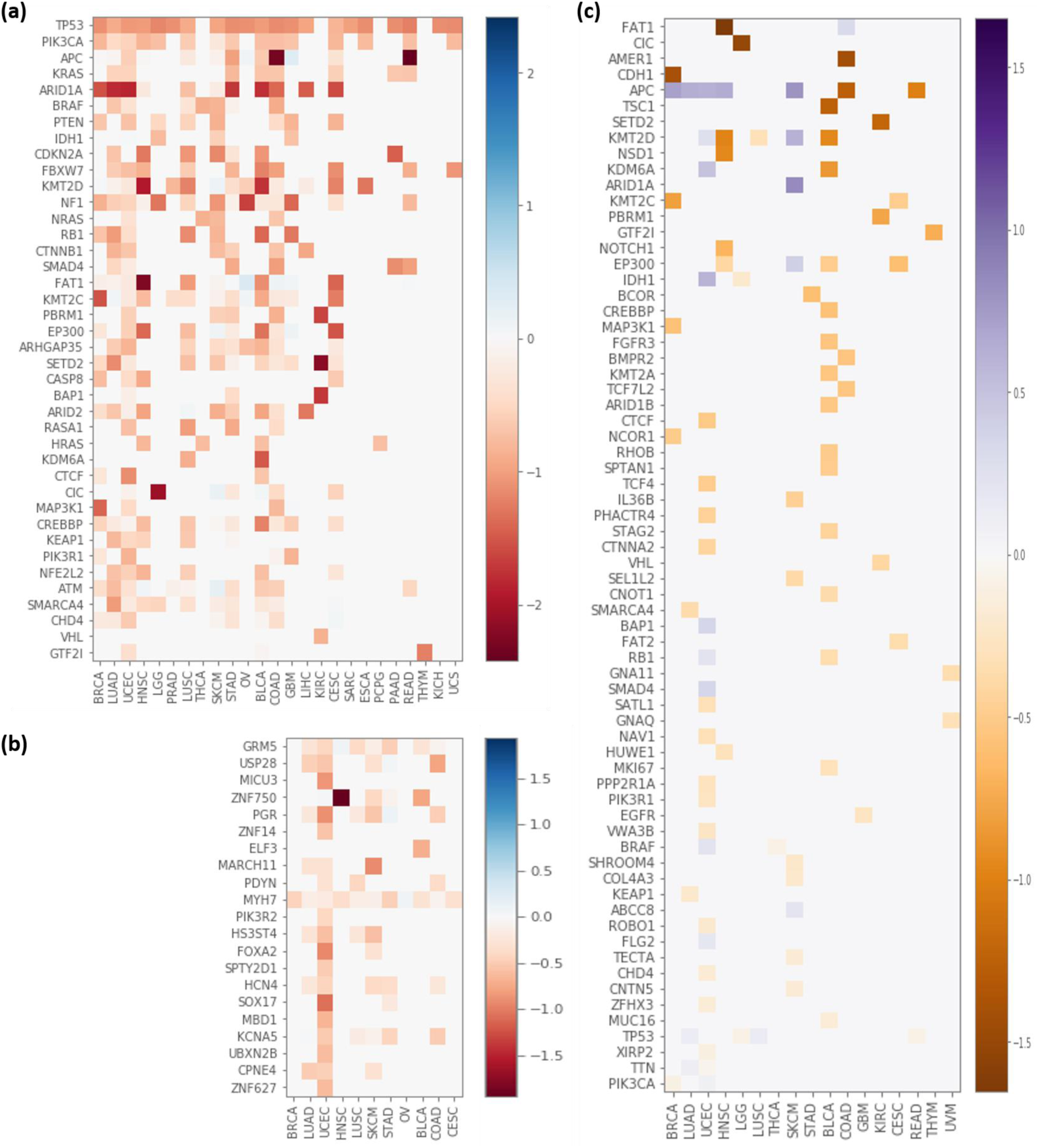
Alteration bias differences across cancer types. The heat maps in (**a**) and (**b**) show the average z-values of the mutation effect scores (compared to the background) across cancer types and genes. More negative values (red) indicate genes that are more biased towards harmful mutations (within the relevant cancer types). We filtered out entries representing the average z-value of less than 15 observed mutations (those are shown as gray 0 values on the map), as averages of too small numbers tend to be noisy and potentially misleading. Note that sparse rows and columns do not imply a lack of alteration bias in the relevant genes or cancer types; they simply indicate small amounts of available data. Only cancer types with at least one remaining entry are shown. (**a**) Top 40 alteration promoting genes, all had been reported as cancer driver genes (see text). (**b**) Highly significant (q-value < 0.001) overlooked genes (i.e. not appearing in Census or CCGD), after keeping only genes with at least 15 observations in at least one cancer type. Genes are sorted by significance. (**c**) The 68 genes found to have significant differences in alteration bias across cancer types. Each value indicates the mean z-value difference between the relevant cancer type to all other cancer types. Negative values (orange) indicate genes that are more damaged in the relevant cancer types (after taking into account the differences in the background models of different cancer types); positive values (purple) indicate genes that are less damaged. Entries (gene and cancer type combinations) that are non-significant (after FDR) are shown as gray 0 values on the map (see Online Methods). Genes are sorted by their largest difference.

ARIDIA, a well-studied cancer driver that belongs to the growing set of drivers found to play a role in chromatin remodeling^33,34^, is a highly significant alteration promoting gene across many cancer types (Fig 4a). However, in Skin Cutaneous Melanoma (SKCM) it is significantly less damaged (Fig. 4c), suggesting that its role in oncogenesis within this cancer type is not as important compared to other cancer types. FAT1 (FAT atypical cadherin 1, also known as FAT tumor suppressor Drosophila homolog) and CIC (capicua transcriptional repressor), both well-known drivers, seem to be particularly dominant in the Head and Neck Squamous Cell Carcinoma (HSNC) and the Brain Lower Grade Glioma (LGG) cancer types, respectively (Fig. 4a and Fig. 4c). FAT1 encodes a cadherin-like protein that binds β-catenin, antagonizing its nuclear localization. Damaging mutations to FAT1 that suppress its binding function thus lead to activation of the Wnt signaling, which is fundamental in tumorigenesis. APC, another tumor suppressor that binds β-catenin, seems especially dominant in the Colon Adenocarcinoma (COAD) and the Rectum Adenocarcinoma (READ) cancer types (Fig. 4a and Fig. 4c). Additional genes that are especially dominant in specific cancer types are: SETD2 in Kidney Renal Clear Cell Carcinoma (KIRC), KMT2C in Breast Invasive Carcinoma (BRCA) and Cervical Squamous Cell Carcinoma and Endocervical Adenocarcinoma (CESC), and GTF2I in Thymoma (THYM); see Fig. 4a and Fig. 4c.

### 2.8 Alteration bias in the healthy human population

So far the presented analysis has focused on somatic mutations in cancer, based on data extracted from TCGA. To further validate our framework, we repeated our analysis on a fundamentally different dataset comprised of germline variants in the healthy human population. Those variants were extracted from ExAC^25^, the largest and most complete contemporary catalogue of genetic variation in the healthy human population.

The full dataset of ExAC contained 10,089,609 variants. We filtered out 1,054,475 low-quality variants (see Online Methods), and among the remaining 9,035,134 variants we found 8,538,742 SNPs. Of these, 4,747,096 were found to be in coding regions, contributing to a final dataset of 4,752,768 gene effects. This dataset was then processed through the same pipeline used for identifying alteration bias in cancer, as described earlier. The effect scores of the variants in each gene were compared against the background distribution derived from the nucleotide substitution frequencies of the observed variants in this dataset.

Unlike in the context of somatic mutations in cancer, where almost all significant results (99%) were found to be alteration promoting genes, here we found the opposite trend. Almost all of the significant results we discovered (97.7%) are alteration rejecting genes. Specifically, we found 6,141 significant alteration rejecting genes, and only 147 significant alteration promoting genes. This finding is not surprising given that normal human evolution is dominated mostly by negative selection, while tumor evolution is mostly dominated by positive selection^35-37^. The full results of all 17,946 analyzed genes are provided in Supplementary Table S1-ExAC.

We tested for enriched GO (Gene Ontology) annotations of biological processes among the alteration rejecting genes with the strongest effect size (in terms of z-value; see Online Methods). The most significant annotations were related to RNA processes (e.g. mRNA metabolic process; FDR q-value = 2.4E-28), RNA splicing (FDR q-value = 9.1E-24) and regulation of gene expression (FDR q-value = 1.3E-23). Additional highly enriched annotations included response to stimulus (FDR q-value = 2.8E-13), regulation of cell cycle (FDR q-value = 7.3E-13) and regulation of translation (FDR q-value = 6.6E-10). We identified many basic cellular processes, mostly those occurring in the nucleus. However, some key processes such as protein trafficking, lipid metabolism and transporting are notable in their absence. The full list of enriched annotations is listed in Supplementary Table S3.

As expected, we find that variants with higher allele frequencies also have larger effect score biases compared to the background distribution (measured by z-values). This correlation is weak (Spearman’s ρ = 0.05) but very significant (p-value < E-300). The tendency of variants with lower allele frequencies to have lower z-values is well apparent in Fig. 5a. This confirms that more harmful variants (with lower z-values) are less likely to become fixed in the population.

**Fig. 5.**
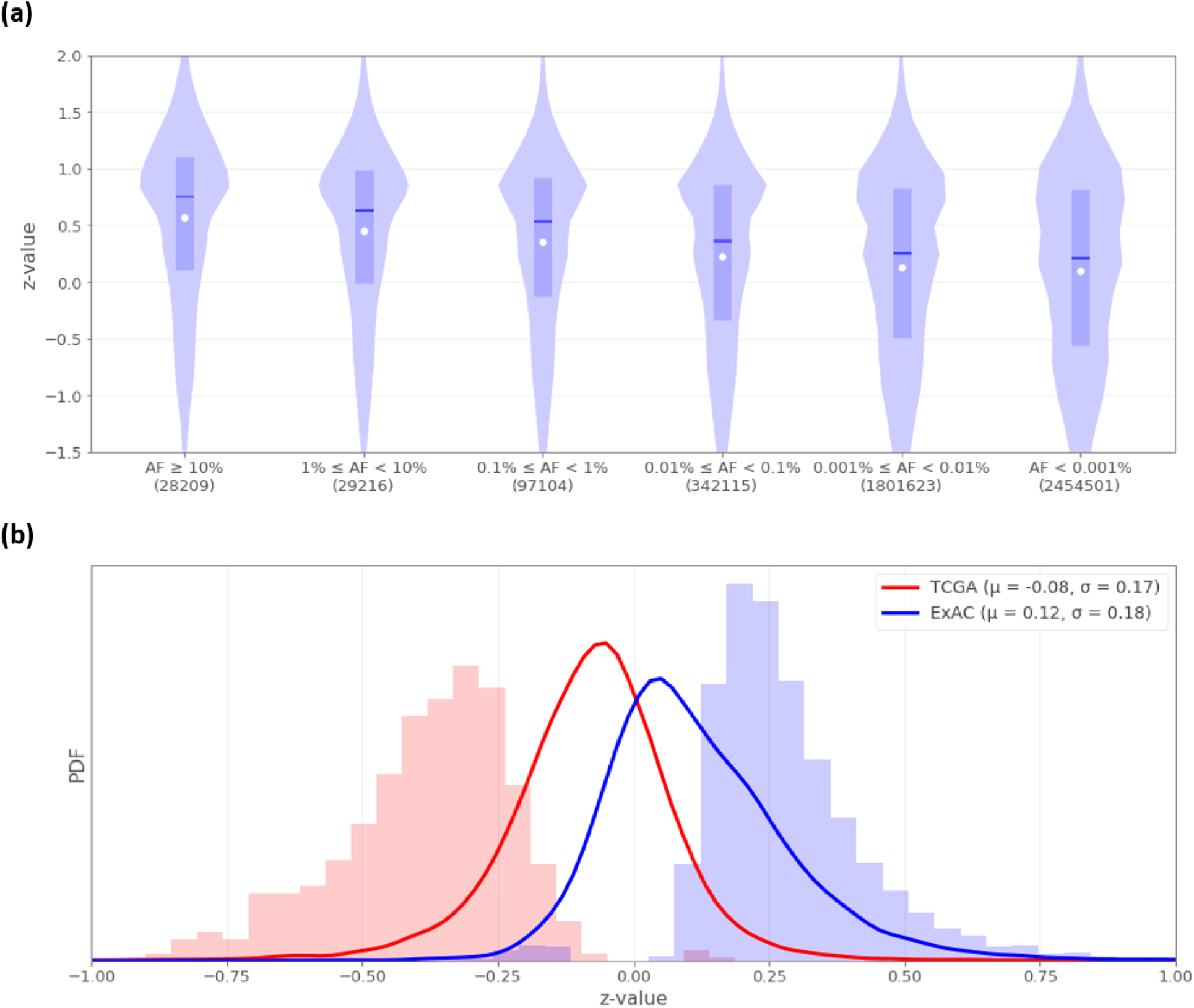
Alteration rejection in the healthy human population. (**a**) Alteration bias (measured by z-value) of germline variants from ExAC across ranges of Allele Frequency (AF). The boxes represent the Q1-Q3 ranges, the middle lines the medians (Q2), and the white dots the means. Since there are ~60k samples in the dataset, the last range (AF < 0.001%) captures only the 2,454,501 effect scores of singleton variants. (**b**) Distribution of alteration bias (measured by mean z-value) of the 17,828 and 17,946 analyzed genes in TCGA (red) and ExAC (blue), respectively. The density plots show the distribution of all analyzed genes, while the shaded histograms only the 599 and 6,288 genes with significant alteration bias in each dataset (comprised of both alteration promoting and alteration rejecting genes in both datasets).

We also found expected correlations between the effect score biases of genes (mean z-values) to other popular scoring techniques that measure evolutionary selection. We report Spearman’s correlation of ρ = -0.4 (p-value < E-300) between the Residual Variation Intolerance Score (RVIS)^38^ to the mean effect score z-values of genes. Similarly, we report Spearman’s correlation of ρ = -0.28 (p-value < E-300) to the Gene Damage Index (GDI)^39^. Both metrics give higher scores to genes that are damaged more than expected, while we give lower scores to such genes; hence the expectation for negative correlation. This further confirms the evolutionary constraints reflected by the effect score biases.

It is also interesting to note a mild overlap between the alteration promoting genes in cancer, found in the analysis of somatic mutations in TCGA, to alteration rejecting genes in the healthy human population, found in the analysis of germline variants in ExAC. Of the 17,313 genes that are shared to both analyses, 584 are significant alteration promoters in cancer, 5,995 are significant alteration rejecters in the human population, and 350 are both. According to random hyper-geometric distribution, we would expect only 202 overlapping genes (x1.73 enrichment, p-value = 1.17E-36). This supports the notion that cancer driver genes, which undergo positive selection during tumor evolution, tend to negative selection during normal evolution.

The notion of opposite trends between cancer to normal human evolutions is strikingly evident in Fig. 5b, which compares the alteration bias in the two datasets. While cancer somatic mutations present a notable bias towards alteration promotion (mean z-value of -0.08 across all genes), germline variants show a similar bias in the opposite direction, towards alteration rejection (mean z-value of 0.12).

To summarize, we developed a general statistical framework and successfully applied it in two fundamentally different domains – cancer genomics and human population genetics. In both territories, the fingerprint of evolution was implied by the functional effect bias of genetic alterations. Our framework is generic enough to unify these two highly different phenomena and to study them with the same analytical tools. We provide the community with catalogues of alteration bias across a wide range of settings. By sharing not only the significant discoveries, but the results of all analyzed genes in each setting, these catalogues can be used as complete references and as baselines for further research.

## 3. Discussion

This work has examined cancer genomics and tumor evolution through the unique perspective of functional alteration bias in protein-coding genes. We developed FABRIC, a machine-learning based framework for detecting alteration bias relying on precise statistical calculations. We discovered 593 significant alteration promoting genes in cancer samples (Table 1), as well as 6,141 significant alteration rejecting genes in the healthy human population. In contrast, we found very few genes showing biases to the opposite directions. The reported list of alteration promoting genes in cancer shows an outstanding overlap with known drivers (Fig. 3), but also reveals many overlooked genes that are plausible driver candidates (Table 2).

Several unique properties distinguish our framework from other methods for detecting cancer driver genes. Most importantly, the underlying assumptions of our framework are minimal. Indeed, the observed effect scores are always compared against a gene-specific background null distribution based on the same prediction algorithm and the same properties of the data (nucleotide substitution frequencies) as the observed mutations. It means that even with a poorly trained prediction algorithm, our framework remains resistant to false discoveries, albeit at the cost of statistical power. If the framework detects that the observed effect scores of a certain gene significantly deviate from its background model, it meant that the somatic mutations within this gene are likely non-randomly distributed.

It is important to remember that the number of observed mutations within a gene has no effect on the strength of its alteration bias. For example, the TPTE gene has one of the highest numbers of observed mutations in TCGA (619 observed SNPs in the coding-region of that gene) and one of the highest mutation densities (0.37 coding-region SNPs per nucleotide, across all samples). Despite that, it shows only a mild alteration bias (average z-value = -0.16) and significance (q-value = 0.005). Unlike conventional methods that look for genes with significantly high mutation densities, our framework treats the number of observations as a baseline for the background model, checking only if the involved mutations are significantly more or less damaging than expected, regardless of their number. This design choice might be strict, but it spares disturbing concerns and doubts that arise in other methods. For example, it has been asserted that many false drivers have been wrongly reported due to the difficulty to construct a proper background model that accounts for highly mutated samples, cancer types and genomic loci^9^. In fact, in many pipelines it is common practice to filter out hyper-mutated samples prior to the searching of driver genes^40,41^, resulting the loss of much of the data. Different choices of normalization techniques and thresholds for hyper-mutated samples may bring very different results, leading to lingering uncertainty. The validity of our framework is not compromised by any of these issues, and hyper-mutated samples can safely remain in the analysis, thanks to our robust background model.

Our method is not only helpful in minimizing false discoveries, but also, we anticipate, in findings new overlooked drivers. Methods that examine mutation densities are based on the premise that cancer driver genes are highly recurrent across patients. However, this assumption may not always apply. Some drivers may appear only among small fraction of the patients, even though they play an important role in tumorigenesis in the rare instances when they do occur. It has been established that tumors are best described by the pathways and protein-protein interaction networks they affect^42,43^. In order for the disease to progress, certain regulatory networks must be disrupted. Some genes, such as TP53, are positioned in the network as central hubs. Other drivers are part of a relatively small network, so their disruption is often necessary for cancer development^44^. Other gene networks, on the other hand, can be larger and more distributed, meaning that damaging any of a large collection of genes would lead to tumorigenesis. In this scenario, genes can be far less recurrent across samples, despite their strong contribution to the disease^45,46^. For example, HOXD11 is a known driver with an important role in tumorigenesis^47^ (which is also captured by our framework; see Supplementary Table S1-TCGA_combined), despite its low recurrence (only 32 observed SNPs in our dataset, across 10,182 samples). Such non-recurrent driver genes may result in mutation densities that are not significantly different from the random baseline, meaning conventional methods would most likely miss them. Our approach is capable of exposing such genes, if the observed mutations substantially disrupt their function.

Within the group of cancer alteration promoting genes, we have focused our discussion on the most well characterized category of tumor functional genes, namely drivers. However, there may also be other functional categories that do not necessarily have a direct role in tumorigenesis per se, but other functions that lead to positive selection in cancer evolution. For example, the alteration of a cancer gene might be positively selected for promoting chemotherapy resistance, after the tumor has already initiated^48^, or for allowing immune system evasion^49^.

Since our framework is based on a signal that is orthogonal to most other methods, namely the content and functional effect of mutations rather than their quantity, the two alternative paradigms can complement and augment each other. Alternative methods that deal with independent elements of the data are ideal for an effective meta-analysis. If a gene is established as a driver by multiple complementary approaches, it provides a very strong evidence for a genuine role in cancer^15,50^.

Among the 593 alteration promoting genes we found, we marked 183 as overlooked driver candidates. We argue that alteration promoting genes are probable tumor actors, and we hope future research will help establishing their role in cancer. In this study we were interested in screening the entire human genome evenhandedly for pursuing driver candidates; validating each discovery is beyond the scope of this work. We provide the full lists of all analyzed human genes, among which are the significant results, in Supplementary Table S1.

In addition to the research of cancer somatic mutations, we also applied our framework on a dataset of germline human variation, comprised of the genetic variants of ~60,000 healthy individuals from ExAC. In this dataset we found a clear bias towards alteration rejection, indicating the strong signature of negative selection in the course of human evolution. This stands out in contrast to the dominancy of positive selection found in cancer. While tumors promote alterations, long-term evolution rejects them (Fig. 5). Our work provides a new angle on the inverse symmetry between tumor evolution to human population evolution. The full list of genes analyzed for their alteration bias in light of human genetic variation, among which are the 6,141 significant alteration rejecting genes and 147 significant alteration promoting genes, is available in Supplementary Table S1-ExAC.

While this work has focused solely on the attempt to uncover genes exhibiting functional alteration bias (mostly in the context of cancer), we anticipate that our approach can have useful adaptations in a much broader context. For example, we propose that a very similar paradigm could advance the discovery of genes in genome wide association studies (GWAS)^51^. In contrast to traditional GWAS methods^52^ that treat all variants equally, we suggest to weigh each variant within the same gene by its functional impact, in the spirit of the approach presented throughout this work. In other words, instead of looking for variants or genes with significantly different allele frequencies between cases and controls (the routine GWAS procedure), one can look for genes with significantly different effect scores between the two groups. This suggested approach can address some major limitations in present-day GWAS^53^. Specifically, it can provide more direct evidence for genotype-phenotype associations than mere variant correlations, pinpointing specific genes. Additionally, a unified gene-centric model can accommodate rare and even de-novo variants. Classical GWAS methods struggle with rare variants^54^, even though they are believed to be the major contributors to most complex phenotypes^55^. We argue that effective aggregation of variants, as demonstrated by our framework, is a potential remedy to these problems.

We advocate for using richer models in genetic studies by explicit modeling of biological processes. In our framework, we modeled the impact of genetic variants with respect to the relevant protein sequences, by extracting features capturing their rich multi-dimensional proteomic context, in order to obtain functional effect predictions. The resulted knowledge-based predictions, and not the raw variants, were the basis of our successful statistical analysis.

Our framework brings together two classes of genetic analysis strategies that are mostly evolving in parallel. The first strategy, exemplified by classical GWAS, aims to find significant associations (e.g. between genotype to phenotype) without explicit modeling of the underlying biological processes. The other strategy attempts to train predictive models, for example to assess the pathogenicity of de-novo mutations (e.g. Polyphen2^56^). Methods of this kind typically rely on explicit modeling of biological principles by relying on prior knowledge. They make extensive use of machine-learning algorithms, and are part of a rapidly evolving field. Machine learning methods are very powerful tools for knowledge-based modeling, as they allow the necessary flexibility for handling with the noise and complexity of real-world data. A major contribution of this work is in providing a schema for using predictive models in the context of rigorous statistical inference, in order to find significant associations. We exploited the knowledge learned by predictive models (assessing the effects of protein alterations), and transformed it into useful biological discoveries (identifying genes with significant alteration bias).

We provide the research and biomedical comminutes with a host of resources and tools. We provide the summary statistics of all analyzed genes in Supplementary Table S1, both in the context of cancer (a combined pan-cancer analysis, and each cancer type separately) and in the context of human population variation. Our framework is also available as an open-source Python project (https://github.com/nadavbra/fabric; see Online Methods) with detailed instructions and documentation, so it can be easily applied in other datasets and adjusted to broader contexts.

## 4. Methods

### 4.1 TCGA data extraction & processing

The dataset of somatic mutations in cancer, used in most of the analyses throughout this work, was extracted from TCGA^19^. Through NIH’s GDC Data Portal^57^, we selected files in the MAF data format that were processed by the MuTect2^58^ workflow for variant aggregation and masking. We selected only open access files, resulting 33 files in our final query (one for each cancer project). The full query URL was:῀

https://portal.gdc.cancer.gov/repository?filters=῀%28op῀%27and῀content῀%28῀%28op῀%27in῀content῀%28field῀%27files.access῀value῀%28῀%27open%29%29%29῀%28op῀%27in῀content῀%28field῀%27files.analysis.workflowtype῀value῀%28῀%27MuTect2%2A20Variant%2A20Aggregation%2A20and%2A20Masking%29%29%29῀%28op῀%27in῀content῀%28field῀%27files.dataformat῀value῀%28῀%27MAF%29%29%29%29%29

In total, these 33 files contained 3,175,929 somatic mutations across 10,182 samples. 2,956,550 of these mutations were SNPs, and 2,235,884 of these SNPs were in coding regions (i.e. substituting a nucleotide within the open reading frame of a protein-coding gene). Each of these coding-region SNPs was assigned effect score(s) for the gene(s) it affected (occasionally it happens that the same mutation affects multiple overlapping genes). According to our pipeline, synonymous and nonsense mutations were automatically assigned with effect scores of 1 and 0 (respectively), while missense mutations were given effect scores by our pre-trained predictor (see details below). Following this logic, we ended up with 2,238,945 effect scores of the processed somatic mutations.

The Python code for parsing and processing the raw data is available as part of our open-source project (https://github.com/nadavbra/fabric).

### 4.2 ExAC data extraction & processing

The dataset of human population variants was extracted from ExAC^25^. We downloaded release1 of the ExAC dataset through their FTP site, obtaining the VCF file at the following URL: ftp://ftp.broadinstitute.org/pub/ExAC/release/release1/ExAC.r1.sites.vep.vcf.gz

The VCF file contained 10,089,609 variants of the human population. Of these, we filtered out 1,054,475 variants with any FILTER flag other than PASS (in the VCF fields). Afterwards, we processed the remaining 9,035,134 variants exactly as we processed the somatic mutations in the cancer dataset (see above). We found 8,538,742 SNPs, 4,747,096 of them in coding regions, contributing to a final dataset of 4,752,768 effect scores.

### 4.3 Constructing gene sequences & annotations

In order to analyze the effects of variants, we needed to construct both the DNA and protein sequences of all human genes. As our machine-learning predictor also required additional features (see details below), we also extracted additional proteomic annotations on top of the protein sequences (e.g. post-translational modifications from UniProt^59^ and protein domains from Pfam^60^). As the TCGA dataset used the GRCh38 version of the human reference genome, while ExAC and ClinVar (see below) used the GRCh37 version (also known as hg19), we constructed the gene sequences and their annotations for both versions.

The genomic coordinates of the human genes were taken from the GENCODE project^61^ (the file gencode.v26.annotation.gtf.gz for GRCh38, and gencode.v19.annotation.gtf.gz for hg19). Using the chromosomal DNA sequences of the human reference genome, downloaded from UCSC^62^, we recovered the DNA sequences of the genes. For GRCh38, we used the reference genome from ftp://hgdownload.cse.ucsc.edu/goldenPath/hg38/chromosomes/, and for hg19 from ftp://hgdownload.cse.ucsc.edu/goldenPath/hg19/chromo/somes/. Importantly, since there is inconsistency in the reference genome of the mitochondria between UCSC and RegSeq/GENCODE (only in the hg19 version), the reference genome sequence for the M chromosome was taken from RefSeq^63^ (NC_012920.1) instead of UCSC. This issue had been resolved in GRCh38. We were interested only in the DNA sequences of the protein-coding regions of genes (annotated as “CDS” in GENCODE).

Next, we downloaded the protein sequences (together with an abundance of proteomic annotations) of all 20,168 reviewed human proteins in UniProt (http://www.uniprot.org/uniprot/?sort=score&desc=&compress=yes&query=organism:%22Homo%20sapiens%20[9606]%22%20AND%20reviewed:yes&fil=&format=xml&force=yes). In order to map between the gene IDs found in GENCODE (Ensembl gene IDs^64^) to the UniProt IDs of the protein sequences, we used the repository of gene names downloaded from genenames.org^65^ (ftp://ftp.ebi.ac.uk/pub/databases/genenames/new/json/non_alt_loci_set.json). genenames.org is a curated online repository of HGNC-approved gene nomenclature. Among other fields, it contains Ensembl and UniProt IDs, as well as official gene symbols. Some genes (identified by a UniProt ID) had multiple matching GENCODE transcripts (possibly due to alternative splicing). In these cases we chose the transcript whose DNA sequence fully matched the amino-acid sequence of the primary UniProt isoform. If no matching transcript was found, we discarded the gene from the analysis. We ended up with 18,052 successfully mapped genes in GRCh38, and 18,053 genes in hg19, each identified with a unique UniProt ID.

Finally, we augmented these ~18k constructed genes with protein domain annotations extracted from Pfam^60^. We downloaded version 30.0 of Pfam for the entire human proteome (ftp://ftp.ebi.ac.uk/pub/databases/Pfam/releases/Pfam30.0/proteomes/9606.tsv.gz). We mapped these domain records onto our genes using the provided UniProt IDs (“seq_id” column in the CSV file).

For each of the final ~18k mapped protein-coding genes we constructed the genomic coordinates of the CDS exons (with respect to the relevant reference genome), DNA and protein sequences, and further annotations from UniProt (e.g. active/binding sites, secondary structure, and post-translational modifications) and Pfam (protein domains). Using the genomic coordinates, we were then able to map variants in the human genome on top of these genes, and determine their protein-level consequences (synonymous, missense or nonsense). For missense variants, we could also extract the relevant proteomic features (reference and alternative amino-acid, and all other protein annotations). This way, we processed all somatic mutations extracted from TCGA, and germline variants from ExAC, and we also processed all possible variants within each gene for constructing the background distributions.

The developed pipeline for merging genomic and proteomic data from various databases into unified gene objects, and deriving the genetic and proteomic effects of genetic variants described at the DNA level, is available as an independent open-source Python library which can be used for general purpose (https://github.com/nadavbra/geneffect).

### 4.4 Genetic variations under consideration and rationale

We restricted our framework to the analysis of SNPs, overlooking other, more complex, types of genetic variations (e.g. indels, copy-number variations and chromosomal rearrangements). Although the vast majority of genetic variations, even in the context of cancer, are SNPs (93% of the somatic mutations in our cancer dataset and 94.5% of the variants in our ExAC-derived dataset were SNPs), it is still likely that this choice entailed the loss of some signal, and with it some significant genes. It is conceivable that our approach underestimates the damage caused to genes, due to the disregard of complex variations such as indels (which may cause major functional alterations like frameshifts).

The main motivation for our choice is its conceptual simplicity and minimal set of required assumptions. Specifically, it is relatively easy to model a null background model of uniformly distributed SNPs in a way that accounts for the observed nucleotide substitution frequencies in each gene (see next section). In contrast, more complex variations such as indels are much harder to model, and it is difficult to properly choose the appropriate mathematical model describing them. Moreover, complicated models require more underlying assumptions, thus jeopardizing the certainty of the obtained results. Still, it is a promising direction for future work to enhance the framework with means to consider more complex variations, thereby adding to its detection power.

Likewise, we restricted the framework to protein-coding genes, and considered the functional effects of genetic variations only within proteins. Even functional effects on splicing that could have been relatively easy to model (e.g. alterations of exon-intron junctions) were ignored, and, all the more so, regulatory elements such as enhancers and promoters. This modeling choice might have caused some devastating genetic variations to be deemed harmless synonymous mutations. Here too, the underlying rationale was simplicity, while perusing a proof of concept for our approach. Our results confirm that the developed method yields powerful results even in its most basic form. Extending the framework to broader contexts of functional effects will definitely be a desirable future improvement. Nevertheless, at least in the context of cancer, protein-coding genes are assumed to be the main actors^3^.

### 4.5 Statistical framework & background model

Our framework uses a pre-trained prediction model for the effect scores of missense variants (see next section for the details of its training). For each variant *v* (in the context of a protein-coding gene) we assign a deterministic effect score *ES(v)* ∈ [0,1] by the following rule: 

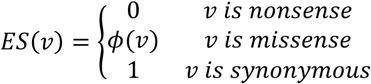

Where *ϕ* denotes the prediction model for missense variants.

In order to construct a background distribution for the effect scores expected at random (Fig. 2), we first consider each single-nucleotide substitution individually. Let *nt*_1_, *nt*_2_ ∈ {*A, C, G, T}, nt*_1_ ≠ *nt*_2_ be two different nucleotides. The background model for the substitution *nt_1_ → nt_2_* in gene *i* is determined by calculating *ES(v)* for all possible substitutions *nt_1_ → nt_2_* within the open-reading frame sequence of the gene. Specifically, let *l_1_,…,l_k_* be all the occurrences of *nt*_1_ within the open-reading frame sequence of the gene. For each *j* ∈ {*1,…,k*}, let us denote by 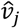 the variant that results upon substituting the occurrence *lj* of nucleotide *nt*_1_ by nucleotide *nt*_2_ within the context of gene *i*. The background distribution for the substitution *nt_1_ → nt_2_* in gene *i*, denoted by 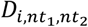, is a uniform distribution over 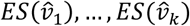 (each chosen with probability 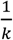).

In order to construct the background distribution *D_i_* for the entire gene *i*, we first calculate the frequencies of the nucleotide substitutions of the observed variants within the gene, denoted 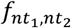 for the observed frequency of the *nt_1_ → nt_2_* substitution. These frequencies satisfy: 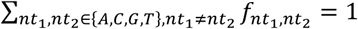. We then take *D_i_* to be a mixture of the twelve 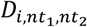 distributions with 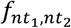 as coefficients (i.e. to sample from *D_i_* one first samples a pair of nucleotides *nt_1_,nt_2_* with probabilities 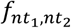 and then samples from 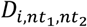).

Let *v_1_,…,v_n_* be the observed variants in gene *i*. We calculate the mean observed score of the gene 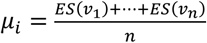 and compare the observed mean *μ_i_* to the background model of the gene, *D_i_*. We do this by calculating the gene’s mean z-value 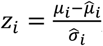, where 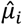 and 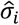 are the mean and standard-deviation of *D_i_*. This is equivalent to calculating the z-value for each variant individually (given by 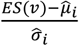) and then averaging them. This value summarizes the overall strength of alteration bias in the variants observed for gene *i*, but it gives no indication of statistical significance. When *z_i_* < 0, gene *i* is potentially alteration promoting, as the observed effect scores are lower than those expected at random, indicating more harmful variants. Similarly, *z_i_ > 0* indicates a potential alteration rejecting gene.

When *z_i_* < 0, we can derive the one-tailed p-value by calculating:

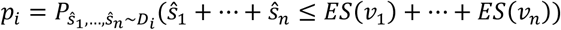

In other words, the p-value is the probability of obtaining scores at least as low as the observed ones, assuming they are independent and identically distributed (i.i.d) according to the background distribution *D_i_*. Similarly, when *z_i_* > 0 we calculate the probability of obtaining scores at least as high as the observed ones. All the reported p-values throughout this work are two-tailed, obtained by multiplying the one-tailed p-values by a factor of 2.

In order to compute the p-values, we need to calculate the distribution of the sum of *n* i.i.d random variables, each with distribution *D_i_*. The distribution of the sum is given by convolving *D_i_* with itself *n* times. To facilitate this computation, we round all the values (both the observed values, and in the background model) to two decimal digits, obtaining 101 distinct bins in the range [0,1]: 0, 0.01, 0.02,…, 0.99, 1. The distribution of the i.i.d sum is then given by 100*n* + 1 bins in the range [0, *n*]. This is equivalent to a precise calculation given that the missense effect score predictor *ϕ* outputs scores in a resolution of *2* decimal places.

As evident from this mathematical formulation, our framework makes no assumptions whatsoever about the validity of the pre-trained prediction model *ϕ*, or the effect-score calculation schema *ES* in general. Even if the scoring function gives arbitrary scores, the calculated p-values are still accurate, and significantly low p-values provide strong evidence against the null hypothesis, namely that the observed variants do not seem to distribute independently and uniformly across the gene. A bad scoring function would undoubtedly diminish the statistical power of the framework, but should not result in false discoveries. The abundance of significant results found in our analysis suggests that our scoring schema is well-designed, and that the machine-learning predictor is well trained. As the background model controls for the prediction model, the number of observed variants and their nucleotide frequencies, the assumptions of our framework are minimal.

### 4.6 Missense and mutation-type analyses

The previous section describes the main form of our framework (referred to as the overall analysis). We have briefly discussed two additional forms, the missense and mutation-type analyses (see Fig. 3). The missense analysis considers only the effect of missense variants, completely ignoring synonymous and nonsense variants. In this form of the framework, all the synonymous and nonsense variants are discarded from both the observed variants and the background variants. Namely, each of 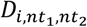 is comprised only of the instances of *nt*_1_ in the gene sequence whose substitutions to *nt*_2_ would result a missense variant. z-values and p-values are calculated for the missense analysis accordingly. Notably, this filtration may substantially diminish the number of analyzed observations, compromising the power of the statistical test accordingly.

The mutation-type analysis, on the other hand is fundamentally different, and does not rely at all on our scoring schema. In this form, only the types of the variants (synonymous, missense or nonsense) are considered, while the heterogeneity of functional effects within missense variants is completely neglected. Mutation-type p-values are calculated by a simple chi-squared test comparing the number of observed variants of each type, denoted *n_syn_, n_mis_, n_non_*, to the numbers that would be expected at 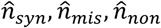, given *n_syn_, n_mis_, n_non_* variants that uniformly distribute across the gene with the observed tendency towards specific nucleotide substitutions (the frequencies 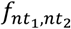 defined earlier). The chi-squared test statistic is then given by

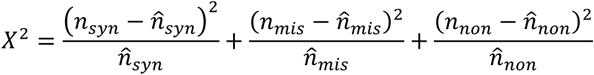
which asymptotically distributes by a *χ^2^* distribution with 2 degrees of freedom under the null hypothesis. In order to meet the asymptotic approximation, we used the accepted rule^66^ requiring that the expected numbers of observations all satisfy 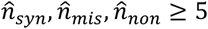 as a prerequisite conditions for the mutation-type analysis. If a gene meets these prerequisite conditions, it obtains a significant two-tailed chi-squared p-value, and it has either more nonsense variants than expected (i.e. 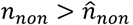) or less synonymous variants than expected (i.e. 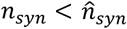) – then it is considered an alteration promoting gene according to the mutation-type analysis. In the combined pan-cancer TCGA dataset, of the 7,674 genes meeting the prerequisite conditions 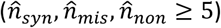, 387 genes were found to be alteration promoting only by their mutation types.

### 4.7 Effect score prediction model

A key component of the framework is a pre-trained machine-learning model for predicting the effects of missense genetic variants on protein function. Given the details of a missense variant, it predicts a numerical effect score between 0 (harmful) to 1 (harmless).

We pre-trained the model on an independent dataset of human genetic variations extracted from CinVar^67^. ClinVar provide a comprehensive catalogue of human genetic variations together with their clinical significance (e.g. pathogenic or benign), as determined by various submitting groups (e.g. OMIM^68^).

We downloaded the full ClinVar dataset from the FTP site at: ftp://ftp.ncbi.nlm.nih.gov/pub/clinvar/tab_delimited/variant_summary.txt.gz. This dataset contained 543,841 variant records, 262,377 of them in version hg19 of the human reference genome. Of these, 215,862 were SNPs, 89,275 of them were missense variants affecting exactly one protein-coding gene. According to ClinVar’s stated clinical significance, we determined whether each variant was pathogenic, benign, or undetermined. Variants that contained the keywords “pathogenic” or “likely pathogenic” were deemed pathogenic; other variants that contained the keywords “benign” or “likely benign” were deemed benign; all other variants were deemed undetermined. Following this logic, of the 89,275 missense records, 22,496 were labeled pathogenic (positives) and 14,512 were labeled benign (negatives). Altogether, we obtained a final dataset of 37,008 records.

Next, we extracted an immense set of features (1,109 in total) for each record, aimed at capturing the rich proteomic context of each missense variant. Those features are discussed in the next section. Once all the features had been extracted, we obtained a dataset of 37,008 records (22,496 and 14,512 of each of the two labels) represented as vectors in a 1,109-dimensional space. We then applied off-the-shelf machine-learning algorithms to train our prediction model, using 3-fold cross-validation to estimate its performance. We chose a Random Forest classifier (implemented by the scikit-learn Python library^69^ with the following hyper-parameters: n_estimators = 100 and min_samples_split = 50. We report the following performance on ClinVar validation sets (average scores of the 3 cross-validation folds): AUC = 90%, F1 = 85.8%, Precision = 86%, Recall = 85.5%, Specificity = 78.4%, Accuracy = 82.7%.

When a machine-learning predictor is used, usually only the predicted label (e.g. pathogenic or benign variant) is of interest, while the exact score given by the prediction model has no significance. Furthermore, the exact scores (usually in the range 0-1) produced by algorithms like Random Forests have no simple interpretable meaning. For our framework we required refined effect scores spanning the entire 0-1 range, preferably with meaningful probabilistic interpretation. To this end, we rescaled the outputs produced by the trained model such that an effect score of *s* ∊ [0,1] would indicate that roughly *s* percentage of the validation-set variants with a similar score were benign (e.g. ~85% of ClinVar’s variants with an effect score of 0.85 were benign). This way, it can be useful to think of a variant with an effect score of 0.85 as having 85% chance of being harmless, although this is by no means guaranteed as we move from ClinVar to other datasets (e.g. TCGA or ExAC), especially considering that ClinVar is highly imbalanced and biased towards having mostly pathogenic variants.

It should be emphasized that the objective of the prediction model is to evaluate the effects of missense variants on the function of the involved genes/proteins, and not on the entire organism (salient human phenotypes). It may seem that our choice of training dataset (ClinVar) violated this declared goal, as the labels extracted from ClinVar (pathogenic vs. benign) were explicitly based on clinical significance, namely at the resolution of the whole organism and not at the molecular level of genes and proteins. However, as detailed in the next chapter, we deliberately included only features that captured the biochemical and biophysical characteristics of the variants at the molecular level, and avoided any feature that would potentially disclose any information of higher-level phenotypes or carry explicit evolutionary context. For example, most functional effect prediction tools (e.g. Polyphen2^56^ and MutationTaster2^70^) use evolutionary conservation of the gene/protein sequence as a primary feature, while we avoided it altogether. Including such features might have boosted our performance on the ClinVar dataset, but, for our goals, it was critical not to give the model any information on the effects of genetic variants at the whole-organism level. Therefore, even though the model was trained to predict clinical outcomes, the features included in its training set revealed information only about the effect of the variant at the molecular level. Therefore, it could only learn to predict molecular functional effects (as a proxy to clinical outcomes).

In this study our goal was not to obtain maximal performance on the ClinVar dataset, but to train a model we could then transfer to a different task – detecting alteration bias. We used the model, that had been pre-trained to predict functional effects at the molecular level, in order to find genes showing significant alteration bias throughout various evolutionary contexts (cancer or human population), even though the prediction model itself had no input regarding evolutionary or clinical information.

The trained model, including an API allowing to invoke it on missense variants, is available as a separate open-source project (https://github.com/nadavbra/firm). All the Python source code for extracting features, training the model on ClinVar, and invoking it, is available within this project. To the best of our knowledge, this is the first tool that aims to predict functional effects at the molecular level and not at the organism level.

### 4.8 Proteomic features used by the prediction model

The developed Random Forest classifier receives missense variants as inputs, each represented as a vector in a 1,109-dimensional space. Those 1,109 numeric values are extracted features about the variants, based on their rich proteomic context. As stated, we included features describing various biochemical and biophysical properties of the affected proteins at the molecular level, but avoided features disclosing information about broader evolutionary or clinical contexts.

The main features included are: i) the location of the variant within the protein sequence, ii) the identities of the reference and alternative amino-acids, iii) the score of the amino-acid substitution under various BLOSUM matrices, iv) an abundance of annotations extracted from UniProt, v) amino-acid scales (i.e. various numeric values assigned to amino-acids, as described elsewhere^71,72^), vi) Pfam domains and Pfam clans. The full specification of all extracted features is available in Supplementary Table S4. The Python source code for extracting the features is also available within our open-source project (https://github.com/nadavbra/firm). The rest of this section briefly describes the features.

We used annotations that appeared in UniProt as “features” within the protein record. These UniProt features include: active site, binding site, (topological) domain, disulfide bond, secondary structure (helix, strand or turn), and various PTMs (e.g. acetylation, (di)methylation, phosphorylation). In order to keep the dimensionality of our feature space manageable, and to allow the detection of meaningful patterns by the machine-learning model, we collapsed some distinct annotations into identical labels. For example, phosphohistidine, phosphoserine, phosphothreonine and phosphotyrosine, all are distinct features in UniProt, were treated simply as “phosphorylation”. Since the amino-acids of the protein sequence were already included as features, we saw no point in repeating the same information multiple times in our feature space. Supplementary Table S5 summarizes all the UniProt annotations we extracted and the labels we assigned them.

Both the UniProt annotations and the amino-acid scales were aggregated in multiple contexts. First, each “hit” of a specific annotation by the residue affected by the missense variant was recorded as a distinct feature, as well as the distance of that residue to the closest residue with that annotation along the protein sequence. In addition, we also included the total count of each annotation along the entire protein sequence, as well as the count of each of the 20 amino-acids. We also included these counts in other contexts with respect to the modified residue (e.g. all the residues within 5 amino-acids of it, or within 10 amino-acids to its right/left). In each of these contexts, we also included summary statistics (average and standard-deviation) of the various amino-acid scales. We also included the scale values of the reference and alternative amino-acids, and their absolute and relative differences.

Using the data extracted from Pfam, we added unique features characterizing the domain closest to the affected residue along the protein sequence. Specifically, we recorded the distance to the closest domain (measured in the number of residues between the modified residue to the domain) and included a binary feature of whether it was a hit or not. If the domain belonged to a common Pfam clan (with at least 25 occurrences in the human proteome), we also recorded the identity of the clan as a feature. Using the HMM models provided by Pfam, we also examined the impact of the amino-acid substitution on the Pfam domain profile (using the HMMER tool^73^). Specifically, we recorded the scores indicating the matching of the protein sequence to the domain profile before and after the substitution. We included these two scores, together with their absolute and relative difference, as 4 additional features. Occasionally it happened that a substitution seemed to have so severely disrupted the HMM model that the protein sequence no longer matched the domain profile after the substitution. This rare occurrence was included as yet another binary feature.

### 4.9 Comparison to external catalogues of cancer driver genes

The Census catalogue^13^ was downloaded from their website at http://cancer.sanger.ac.uk/census/, in CSV format. We used the “Gene Symbol” and “Role in Cancer” columns in order to determine the cancer role, according to Census, of each of the analyzed genes (identified by its official gene symbol). These possible cancer roles are: i) TSG (Tumor Suppressor Gene), ii) oncogene and iii) fusion. In Fig. 3 we considered a gene to be a “driver” according to Census if it was annotated with any of these roles; Supplementary Table S2 also includes more refined comparisons.

The CCGD dataset^18^ was downloaded from their website (http://ccgd-starrlab.oit.umn.edu/download.php) in CSV format (http://ccgd-starrlab.oit.umn.edu/dump.php). We used the “Human Symbol” and “Studies” columns in order to determine the number of studies supporting each human gene as a driver. In Fig. 3 we compared the list of significant alteration promoting genes to genes supported by at least 10 studies according to CCGD; in Supplementary Table S2 other thresholds are also attempted.

The list of 106 genes recognized as drivers by the MutSig suite was downloaded from Supplementary Table 2 of the published paper^31^. We took the 114 genes that had a significant pan-cancer q-value (according to column BE in the obtained Excel file). Of these 114 gene symbols, 106 had a corresponding gene in our analyzed dataset.

Testing for overlapping between the list of significant alteration promoting genes in cancer to each of these three external lists was carried out with Fisher’s exact test (given the null hypothesis of independence, i.e. hypergeometric distribution).

### 4.10 Highlighting significant differences among cancer types

In order to measure differences in alteration bias among cancer types (Fig. 4c), we collected all the observed SNPs within the coding region of each gene, and for each of the 33 cancer types we split these mutations into two disjoint groups: those that came from patients with that cancer type versus all the others (i.e. from patients with any of the other 32 cancer types). Based on the nucleotide substitution frequencies of the mutations within each group, we calculated a null background model for the effect scores within each of the two groups separately (see the “Statistical framework & background model” section). Based on the effect scores and the background model of each of the two groups, we calculated z-values for the mutations within each group. As the z-values directly measure alteration bias (after accounting for the different background models), they can be compared between the two groups. Indeed, we calculated the differences between the z-values and obtained confidence intervals for these differences (i.e. between the z-values of the particular cancer type to the z-values of all other cancer types, within the same gene). The calculation of confidence intervals was carried out with a pooled two-sided t-test, using the implementation of the statsmodels Python library^74^: statsmodels.stats.weightstats.CompareMeans.tconfint_di ff. A p-value for the difference was calculated using SciPy’s^75^ two-sided t-test: scipy.stats.ttest_ind.

A combination of gene and cancer-type was considered significantly different (compared to all other cancer types, with respect to the same gene), if: 1) there were at least 25 relevant observed mutation of this combination, and 2) it had a significant p-value after FDR. The confidence intervals (uncorrected for multiple testing) and q-values (FDR corrected) of all gene and cancer-type combinations are reported in Supplementary Table S1-TCGA_diff. Of these, 92 combinations were significant (q-value ≤ 0.05). These 92 combinations occurred across 68 genes and 17 cancer types, and are presented in Fig. 4c. The heat map values were taken to be the most moderate values of the confidence intervals (i.e. the lower bound for purple-colored positive values, and the upper bound for orange-colored negative values).

### 4.11 Enrichment testing for gene ontology annotations

In order to test for significant Gene Ontology (GO) annotations^76^ enriched within the genes that most strongly reject alteration in the context of human evolution, we prepared a list of genes with the strongest effect size (in terms of z-value). To avoid statistical noise of small numbers, we only considered the 14,343 genes with at least 100 observations (i.e. at least 100 relevant unique variants in the final dataset extracted from ExAC). Of these, we compiled a list of 227 genes with a very strong effect size, defined by z-value ≥ 0.5. We checked for enriched annotations within this list using the GOrilla tool^77^, available at: http://cbl-gorilla.cs.technion.ac.il/. We used “Homo sapiens” as the organism, and selected the “Two unranked lists of genes (target and background lists)” option. We put the list of 227 strong alteration rejecting genes as the “Target set”, and all 14,343 considered genes as the “Background set”. Of the 14,343 and 227 entered UniProt IDs, GOrilla recognized 13,901 and 225, respectively. We selected only “Process” ontologies. The resulted list of enriched annotations is available in Supplementary Table S3.

### 4.12 Source code availability

We provide all of our source code (written in Python) as open source projects in GitHub. For better usability, we created three different repositories:

1. https://github.com/nadavbra/geneffect
2. https://github.com/nadavbra/firm
3. https://github.com/nadavbra/fabric

Our code relies on the following Python packages: IPython^78^, Cython^79^, NumPy^80^, SciPy^80^, Matplotlib^81^, Pandas^82^, scikit-learn^69^, StatsModels^74^, Biopython^83^ and interval_tree (https://github.com/moonso/intervaltree). The first repository contains a Python package we wrote for combining genetic and proteomic data into unified gene objects, and for interpreting the effects of genetic variants on proteins (see the “Constructing gene sequences & annotations” section). This package is appropriate for general use.

The second repository contains our trained model for assigning effect scores to missense mutations (see the “Effect score prediction model” section). It contains all the necessary code and data files for using the trained model, and for training it from scratch (for reproducibility).

The third repository (which depends on the first two) contains the FABRIC framework for detecting genes showing alteration bias from input data of observed variants. It can be used for detecting alteration bias in cancer (using data from TCGA) or in the healthy human population (using data from ExAC) as demonstrated in this work, and it can also be used on any other dataset from which genetic variants can be derived. In addition to providing FABRIC as a general-purpose tool, this repository also contains all the source code necessary to replicate all the analyses, tables and figures in this work.

## Acknowledgements

We would like to thank Or Zuk from the Department of Statistics at the Hebrew University of Jerusalem for his very helpful advice and insight which have contributed to the statistical foundation of this work.

We would also like to thank Roni Rasnic from the School of Computer Science and Engineering at the Hebrew University of Jerusalem for her valuable review of the manuscript.

## Supplementary Information

**Supplementary Table 1:** Full results (per gene) of all the analyses in this work (based on TCGA and ExAC datasets).

**Supplementary Table 2:** The extent of overlap between each pair of methods and resources for identifying alteration promoting or driver cancer genes (source material for Fig. 3).

**Supplementary Table 3:** GOrilla process annotations enriched within the strongest alteration rejecting genes in ExAC.

**Supplementary Table 4:** Proteomic features used by the prediction model.

**Supplementary Table 5:** Tracks extracted from UniProt, as part of the proteomic features.

